# Physiological costs of prospecting in a resident cooperative breeder, the Florida scrub-jay

**DOI:** 10.1101/2023.07.10.546769

**Authors:** Young Ha Suh, Conor Taff, Reed Bowman, John W Fitzpatrick

## Abstract

Prospecting is an understudied yet pivotal information-gathering process often preceding natal dispersal. While prospecting may enable individuals to optimise dispersal outcomes and obtain high quality territories, it is also likely to incur costs stemming from energy expenditure and predation risks. This trade-off may drive individual differences in prospecting effort. We tested for evidence of costs of prospecting behaviour in a wild population of Florida scrub-jays, *Aphelocoma coerulescens*, which prospect as nonbreeding helpers. Using random sampling across all helpers, we compared prospecting effort—approximated by frequency, distance, and activity level—with body mass changes and oxidative stress levels. We tested if prospecting incurred costs and if early-life body condition predicted subsequent prospecting effort. Prospecting frequency was positively linked to oxidative damage but not to any loss in body mass during the breeding season, suggesting that extra-territorial movement costs manifest unevenly. Early-stage body condition did not affect subsequent prospecting effort across a large set of helpers, but early-stage body condition and morphometric measures did correlate with subsequent oxidative status of those sampled. Our results suggest that prospecting movement carries some physiological costs, perhaps contributing to individual differences in prospecting. This research highlights how body condition plays a role in trade-offs between information gathering movement and physiological costs of movement itself, ultimately providing insight on the evolution of prospecting in social species.

**HIGHLIGHTS:**

- Prospecting prior to breeding enables informed dispersal but incurs cost
- We tested whether prospecting by nonbreeding Florida scrub-jays results in physiological costs or varies with initial condition
- We measured oxidative status using assays testing antioxidant capacity and oxidative damage
- Frequent prospecting was linked to subsequent oxidative damage
- Early-stage body condition and wing length affected subsequent oxidative status

## INTRODUCTION

Natal dispersal is a fundamental life-history event that underlies gene flow and drives population dynamics (Clobert et al., 2012). While often regarded as a random process, natal dispersal – particularly in vertebrates – has been shown to depend on condition and surrounding social and environmental information (Clobert et al., 2009; Delgado et al., 2014; Ponchon et al., 2021). Increasing evidence suggests that animals gather information that aids in optimised dispersal decisions, helping individuals attain high-quality territories (Doligez et al., 2004; Pärt et al., 2011; Schjørring, 2002), ultimately contributing to lifetime fitness (Baines & McCauley, 2018; Bowler & Benton, 2005; Stamps, 2006). Despite its potential fitness benefits, this process of gathering information around one’s surroundings prior to dispersal, termed “prospecting” (Reed et al., 1999), has been shown to involve costs associated with extra-territorial movements. Such costs involve energetics of flight (Butler, 2016), increased exposure to predators (Johnson et al., 2009), and aggression from competitors (Kingma et al., 2016; Smith, 1978). In cooperatively breeding species, such costs are hypothesised to drive delayed dispersal as natal territories provide safe havens for offspring (Koenig et al., 1992; Kokko & Ekman, 2002; Russell, 2001; Tanaka et al., 2016; Tarwater & Brawn, 2010; Woolfenden & Fitzpatrick, 1984). Cooperative breeders have an additional potential trade-off, because prospecting behaviour might occur at the expense of group-based activities such as parental care and territorial defense that can advance both direct and indirect fitness (Barve et al., 2020; Young et al., 2005). Therefore, identifying costs of prospecting is important for improving our understanding of the trade-offs involved in natal dispersal and evolution of group living in vertebrates.

The main constraints to prospecting presumably arise from physiological costs and limits related to body condition or self-preservation (Delgado et al., 2014). As prospectors navigate unfamiliar spaces—potentially evading predators and avoiding aggression from territory-holding conspecifics—increased activity can incur physiological costs that reduce body condition, which can affect survival or reproduction in the long run (Schuett et al., 2012). Indeed, active prospectors often decline in body condition over time, as seen in prospecting meerkats, *Suricata suricatta*, and southern pied babblers, *Turdoides bicolor*, which lose body mass once they move away from their natal group (Maag et al., 2019; Ridley et al., 2008; Young & Monfort, 2009). Even in Seychelles warblers, *Acrocephalus sechellensis*, that live in a predator-free environment, prospectors exhibited reduced body mass compared to non-prospectors (Kingma et al., 2016). In facing these challenges, prospectors either require the capacity to cope with physiological costs or need to be in good condition prior to prospecting. While testing causality in the field is difficult to achieve, indicators of body condition could help elucidate the relationship between physiology and prospecting.

As a component of body condition and a mediator of life-history trade-offs, oxidative metabolism is one mechanism that can explain the link between physiological costs and prospecting behaviour. As a consequence of cellular metabolism, reactive oxygen species (ROS) are unavoidably produced and oxidative stress occurs when organisms cannot effectively mitigate the damaging effects of ROS (Finkel & Holbrook, 2000; Monaghan et al., 2009). With prospecting movement likely involving cellular processes, oxidative metabolism may also mediate trade-offs in the process of informed dispersal (Costantini et al., 2008; Monaghan et al., 2009). Another life-history trait associated with increased energetic and metabolic demands, reproductive effort has been linked to oxidative damage (e.g., Nilsson 2002; Losdat et al. 2011; Heiss and Schoech 2012). Extra-territorial movements may also cause oxidative damage by elevating chronic stress that produces glucocorticoids, which are catalysts for cellular oxidation (Costantini, 2014b; Costantini et al., 2011). Studies in meerkats demonstrated that prospecting males and dispersing females exhibit elevated fecal glucocorticoid metabolite levels compared to residents (Maag et al., 2019; Young & Monfort, 2009), substantiating that the extra movement causes oxidative damage. Testing whether prospecting movement correlates with oxidative damage can reveal how prospecting may indeed be constrained by physiological trade-offs.

Here, we investigated whether prospecting has physiological costs in a resident cooperative breeder, the Florida scrub-jay, *Aphelocoma coerulescens*. We used field measures of body condition and assays of oxidative status alongside estimates of prospecting effort. Previous studies on Florida scrub-jays have demonstrated mass changes (Brand & Bowman, 2012; Cucco & Bowman, 2018) as well as oxidative damage (Heiss & Schoech, 2012) in relation to breeding season and reproductive effort. However, the relationship between such proxies of body condition and prospecting effort remains unknown owing to limited information on prospecting itself. Our study is one of the first to test how prospecting movement correlates to body mass changes and oxidative damage.

We hypothesised that prospecting entails physiological costs in Florida scrub-jays, potentially constraining prospecting effort and leading to individual differences. In one scenario, prospecting efforts may lower body condition over time so individuals that prospect more will have higher measured costs. In another scenario, initial body condition may constrain subsequent prospecting effort because individuals in good condition can afford these costs while those in poor condition cannot spare additional costs of movement (condition-dependence). While these scenarios are not mutually exclusive, and causality is difficult to confirm in the absence of controlled experiments, either pattern would suggest existence of a physiological trade-off between prospecting behaviour and body condition. We sought to understand the physiological constraints of prospecting by first asking whether prospecting is physiologically demanding by testing whether prospecting movement positively correlated with weekly mass loss and/or oxidative stress measures. Next, we assessed whether prospecting is condition-dependent by testing if initial body condition determined prospecting effort throughout the breeding season. Last, we evaluated whether early-stage body condition affects future oxidative status by comparing morphometrics and body condition of nestlings and juveniles to their oxidative condition as yearlings.

## METHODS

### Study system and species

We conducted our study on an intensively monitored colour-banded population of Florida scrub-jays at Archbold Biological Station (hereafter Archbold) in central Florida (27.10°N, 81.21°W) during the 2019 breeding season. The Florida scrub-jay is a cooperatively breeding corvid inhabiting xeric oak scrub habitat in which groups defend territories year-round. Long-term study efforts at Archbold include mapping of territory boundaries by using playbacks and behavioral observations throughout the breeding season, when territorial displays are most intense (Fitzpatrick & Bowman, 2016). In this cooperative breeder, offspring delay dispersal for at least one year on the natal territory as nonbreeding helpers that participate in territory defense, sentinel behaviour, and co-rearing of young (Woolfenden & Fitzpatrick, 1984). Throughout their non-breeding period, helpers can also be seen outside the natal territory boundary during what have been described as “prospecting forays” where they visit other territories but eventually return home at the end of the day (Woolfenden & Fitzpatrick, 1984). Most prospecting forays occur just before the start of the breeding season, around February, but reduce in frequency as the breeding season progresses and nestlings and fledglings benefit from increased provisioning by all group members. In addition, prospectors are often observed at territory boundaries where the presence of non-natal helpers can precipitate a defensive response from the breeding pair or resident group. Territorial defense involves loud vocalizations and exaggerated flight displays, which further attract other birds in the area and result in temporary aggregations (Woolfenden and Fitzpatrick 1984; pers. obs.).

### Collecting prospecting movement

We collected prospecting data through “aggregation sampling” in which we recorded identities of all Florida scrub-jays seen at previously defined sampling points (details and validation in Tringali et al. 2020). To ensure maximum coverage of the study tract, sampling points were non-randomly stratified given past year’s territory boundaries and placed at least 200 m apart (Fig. 1). Each sampling point was visited twice a week and the visiting order was randomized through a random number generator. We played pre-recorded territorial calls of non-extant birds for a total of 2.5 min in each cardinal direction with 30 second breaks in between. In addition to aggregation sampling, we opportunistically recorded aggregations of birds located during the spring breeding season. All identities detected at a sampling point were recorded in the field using Survey123 (Esri, Redlands, CA, USA). To quantify prospecting movement, we calculated each identified individual’s instances seen at a survey point (“frequency”), maximum distance away from the centroid of the individual’s natal territory (“distance”), and proportion seen farther than twice the width of an average territory (592 meters; “activity level”). Aggregation sampling occurred for 8 weeks from mid-February to early April, when breeding season reached its peak.

**Figure 1.**
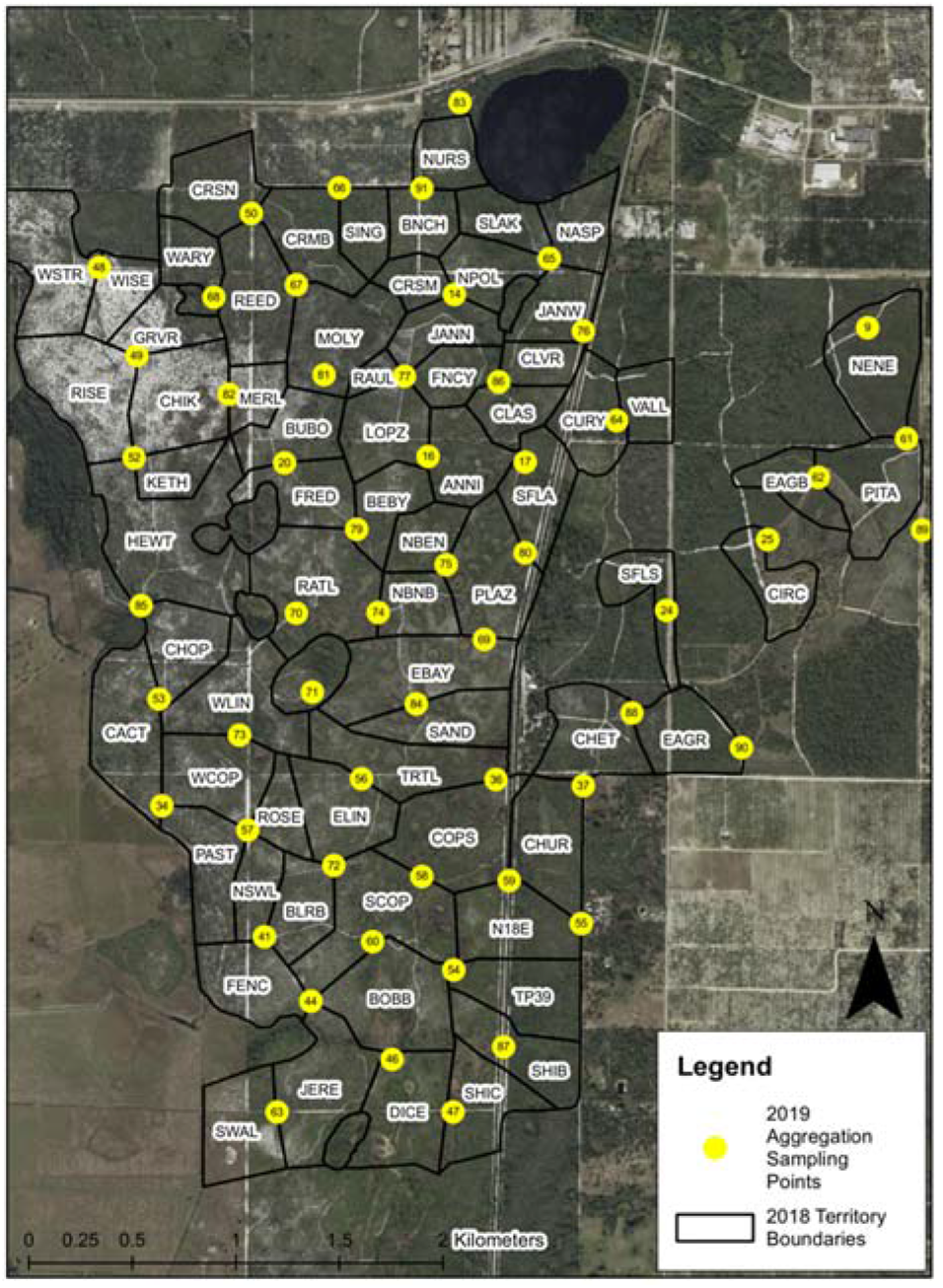
Aggregation sampling points throughout the study tract. Yellow circles indicate 2019 sampling points and black lines indicate territory boundaries (territory names indicated in 4-letter codes) as mapped in 2018.

### Sampling blood and morphometrics

As part of the long-term demography study, all colour-banded Florida scrub-jays are measured and sampled at 11 days post-hatching (nestling) and at around 85 days post-fledging (juvenile). In addition to this effort, we collected morphometric measures and blood samples at approximately 12 months old (yearling) for 28 nonbreeding individuals, all of which were exhibiting the typical range of prospecting behaviours to a greater or lesser extent. Juveniles and yearlings were captured using Potter traps baited with peanuts. We monitored all baited traps and extracted birds immediately after capture (< 1 minute). Morphometric measures included tarsus length (in mm), length of the 7^th^ primary feather (in mm), and body mass (in g) for all age groups. For blood samples, we collected 200–400 µL of blood (<1 % of body mass regardless of age group) via brachial venipuncture and heparinized capillary tubes. All blood samples were taken within 2 minutes of capture for juveniles and yearlings. We centrifuged the capillary tubes at 3,000 rpm for 5 minutes to separate the plasma from red blood cells. We froze the samples in −80°C before transporting them back to Cornell University for subsequent analyses.

### Measuring body mass changes

We used a digital scale (NV511 Precision Balance, Ohaus Corporation, Parsippany, NJ; repeatability of 0.1 g with < 1 second stabilization time) equipped with a feeding tray to obtain body mass measurements of yearlings at the start of the breeding season in 2019. We trained birds to stand on the feeding tray using peanuts as a small food reward. We recorded weighing trials with a video recorder on a tripod panning the scale LED and scored for accurate measurements (Fig. 2). We waited until each individual was on the scale at least three times if possible (degree of habituation varied between individuals) and averaged the obtained weights of each trial (SD ± 0.4g). Birds usually only ate one peanut (ca. 0.1g) at a time and cached all subsequently obtained peanuts, hence any food rewards had negligible effects on body mass data. Because Florida scrub-jays exhibit diurnal mass gain (Cucco & Bowman, 2018), we measured weights only between 7:00 and 10:00 a.m. For each individual, we calculated body mass differences throughout the breeding season (final value minus initial value). We excluded birds with only one weight measure, resulting in 44 individuals that include both yearlings and older birds (*N_yearlings_* = 26, *N_older_* = 18).

**Figure 2.**
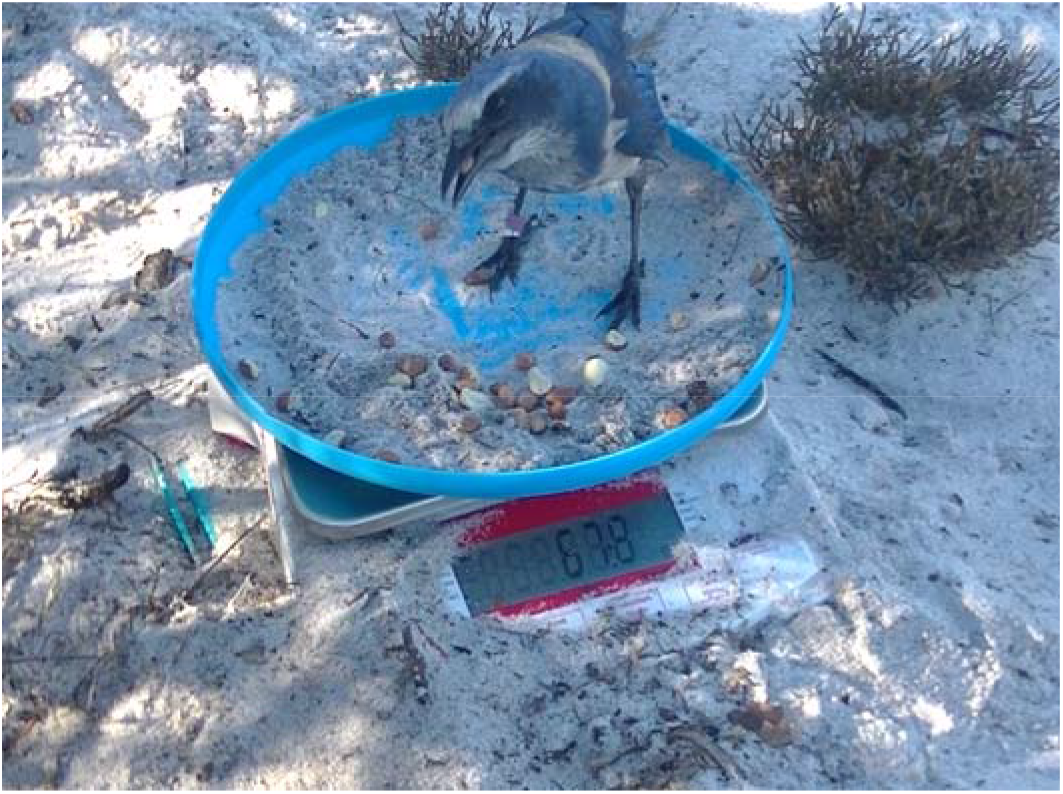
Florida scrub-jay on the scale, showing the digital mass reading and individual colour bands. The scale was partially buried in sand to prevent neophobic responses (avoidance, pecking, etc.).

### Measuring oxidative stress

We measured oxidative status in blood plasma using two commercially available kits that assess total antioxidant capacity (OXY) and reactive oxygen metabolites (ROMs) (Diacron International, Grosseto, Italy). Both assays have been widely used in birds and other animals (Costantini, 2014a).

The OXY-adsorbent test (OXY-Ads) measures total non-enzymatic antioxidant capacity of plasma. We closely followed the kit directions except that we adjusted sample volumes for use on a microplate by diluting 1μL plasma with 99μL distilled water (Treidel et al., 2013). We incubated the plate at 37°C for 10 minutes with a 200μL aliquot of hypochlorous acid solution, which was then added with 5μL of chromogen. We read absorbance at 505nm wavelength using a BioTek Synergy HTX microplate reader.

The d-ROMs test measures the derivative ROMs and indicates the amount of oxidative damage from ROS (Costantini, 2016). We followed manufacturer’s protocols but reduced reaction volumes to fit a 96-well plate by diluting 5μL of plasma into 200μL acetate buffer mixed with 2μL chromogen. We incubated the plate at 37°C for 75 minutes then measured absorbance at 505nm wavelength using a BioTek Synergy HTX microplate reader. For both assays, we ran all samples in duplicate and we used reference standards and blanks for calibration as described in the kit manuals.

We ran both oxidative assays for plasma collected across three age groups and obtained an average value for each individual (first-year mean of 3 measures) and individual slope of change (final measure/initial measure). Of 28 individuals with plasma samples, 23 individuals had complete time series (3 measures) while 5 individuals missed one or more because of missed nestling sampling (nest not being detected by day 11), disappearance during independent sampling, or being trap-shy.

### Statistical analysis

We first tested if prospecting movement corresponded to mass loss and oxidative stress measures. We used linear mixed models (LMMs) to test if prospecting indices—frequency, maximum distance, and activity level—affected oxidative status (first-year average OXY-Ads, OXY-Ads changes, first-year average d-ROMs, and d-ROMs changes) and body mass changes throughout the breeding season. We included sex as a predictor in all our models to account for sex-differences in dispersal in this species (Fitzpatrick et al., 1999; Suh et al., 2020).

Next, we tested if early-life body condition corresponded to subsequent prospecting effort using LMMs and generalized linear mixed models (GLMMs) with a larger pool of individuals (*N_males_* = 23, *N_females_* = 21) that had both early-life morphometric measures and prospecting indices. With LMMs, we tested how body condition as both nestlings and juveniles affected prospecting frequency and maximum distance, and with a GLMM, we tested how the same predictors affected activity level as a binomial response (*N_active_* = 33, *N_sedentary_* = 11). We calculated an index for body condition using residuals from an ordinary least squares linear regression of body mass against tarsus as a linear body size indicator (Pärt, 1990). We selected tarsus as the body size indicator given its weak correlation with other body measures such as wing length and how it remains constant throughout an individual’s lifetime (Green, 2001; supplementary material Table S1). We calculated residuals by sex and added age as a predictor (yearling vs. older helper, *N_yearlings_* = 26, *N_older_* = 18) to account for potential age effects.

Last, we tested whether early-stage body condition corresponded to oxidative status as yearlings. Using LMMs, we tested if body condition as a nestling or juvenile related to OXY-Ads or d-ROMs measured as a yearling. Since incorporating additional measures helps provide a more comprehensive index of body condition (Labocha & Hayes, 2012; Waye & Mason, 2008), we included length of the 7^th^ primary feather in both early-age stages in the models as 7^th^ primary measures were not significantly correlated with body condition (Spearman’s rank correlation all *P* > 0.1; Supplementary material Table S1).

We conducted all analyses in R version 4.0.2 (R Core Team, 2021). To test for repeatability, we used the “rptR” package v. 0.9.22 (Stoffel et al., 2017) assuming a Gaussian distribution and using 1000 parametric bootstraps to evaluate uncertainty. For LMMs and GLMMs, we used the “lme4” package v. 1.1-23 (Bates et al., 2015) and “lmerTest” package v. 3.1-3 (Kuznetsova et al., 2017) and reported parameter estimates, standard error, z- and p-values of the full models, along with R^2^ values using the “MuMIn” package v. 1.43.17 (Barton, 2020). To facilitate interpretation, we scaled all predictor variables by centering the parameters on 0 and dividing them by their standard deviations. We included group identity as a random effect in all models but later removed it in certain models, where its variance component equaled zero (i.e., prospecting effects on weight change, early-life morphometric effects on yearling OXY-Ads). We tested all model assumptions in the “DHARMa” package v. 0.4.4 (Hartig, 2022) and checked for multicollinearity using variable inflation factors (all below 1.9). We created all figures with the R packages “ggplot2” v. 3.3.2 (Wickham, 2016) and “ggeffects” v. 2.8.10 (Lüdecke, 2018).

## RESULTS

Florida scrub-jay helpers were detected prospecting with an average frequency of X ± SE = 9.6 ± 0.4 encounters during the 8-week sampling period. Average maximum distance away from the individuals’ natal territory centroid was 844.6 ± 55.1 m, or about 1.4 times an average territory width (*N* = 202; supplementary material Table S2). Florida scrub-jays experienced remarkably small body mass changes over the breeding season (mean loss −0.03 ± 0.01 g). When split by sex and age, older female helpers were the only group that slightly gained body mass on average (0.8 ± 1.2 g; supplementary material Table S3) whereas yearling helpers lost the most body mass on average (−1.1 ± 0.7). OXY-Ads levels increased with age group (*P* < 0.001; Fig. 3A) but age-adjusted repeatability was not significant (*R* = 0.07, SE = 0.1, CI = [0,0.32], *P* = 0.28). On the contrary, d-ROMs did not change significantly with age group (*P* = 0.11; Fig. 3B) and similarly, age-adjusted repeatability was not significant (*R* = 0.05, SE = 0.10, CI = [0,0.32], *P* = 0.37).

**Figure 3.**
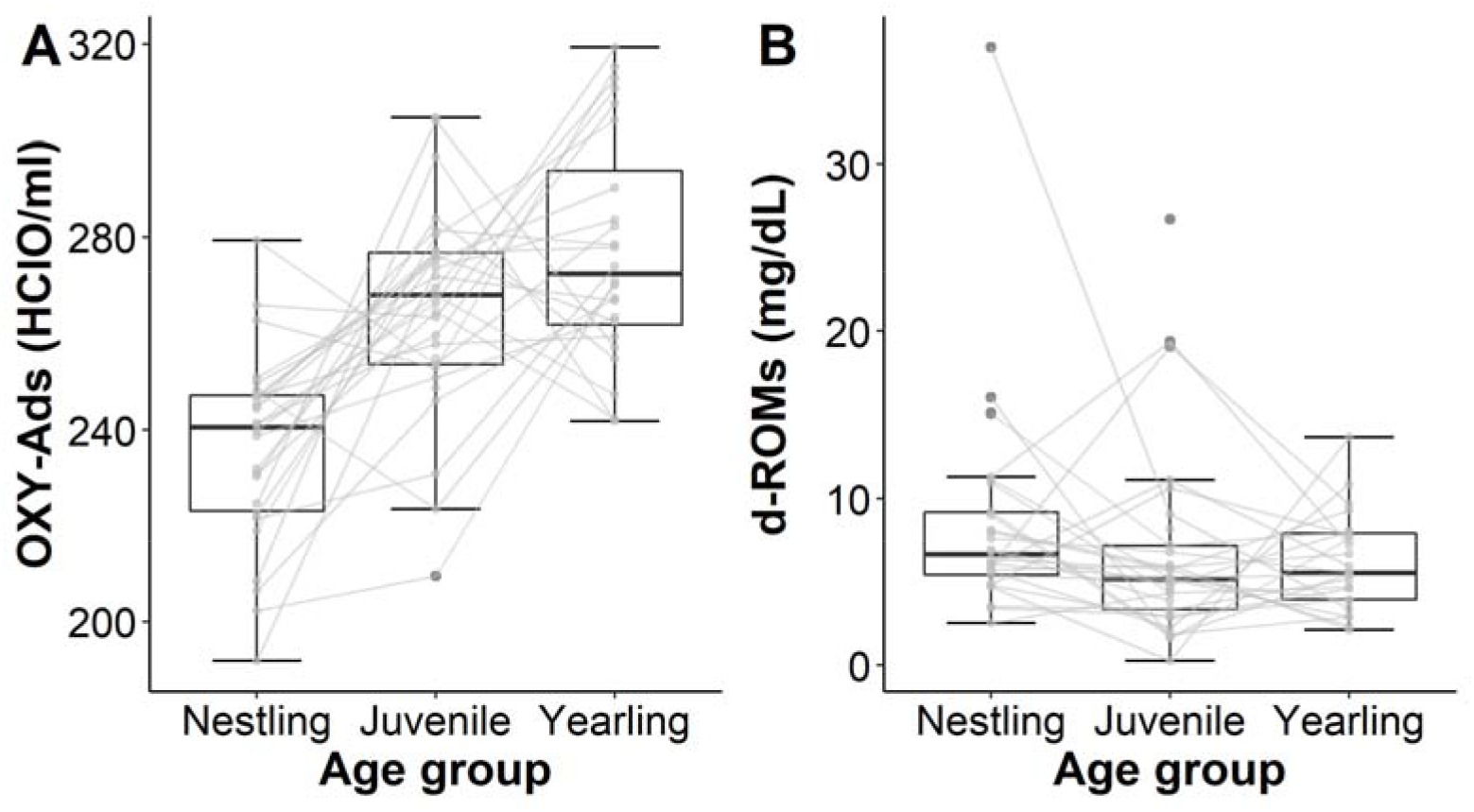
Measures of OXY-ads (A) and d-ROMs (B) by age group with individual age-based measures overlayed and connected in gray. Hinges of boxplots correspond to the first quartile, median (thick line), and third quartile and whiskers indicated 1.5 times inter-quartile range.

### Is prospecting costly?

While prospecting frequency did not significantly vary with OXY-Ads related measures (*N* = 28; β = −3.46, SE = 3.63, *z* = −9.50, *P* = 0.36; Fig. 4A), it significantly varied with first-year average d-ROMs levels, with more frequently detected birds having higher oxidative damage (*N* = 28; β = 2.30, SE = 1.07, *z* = 2.15, *P* = 0.04; Fig. 4B). Neither prospecting measures nor sex affected first-year averages of OXY-Ads or changes in either oxidative status (*P* > 0.2; Table 1). Last, prospecting indices did not influence body mass changes (*P* > 0.3; Table 1).

**Figure 4.**
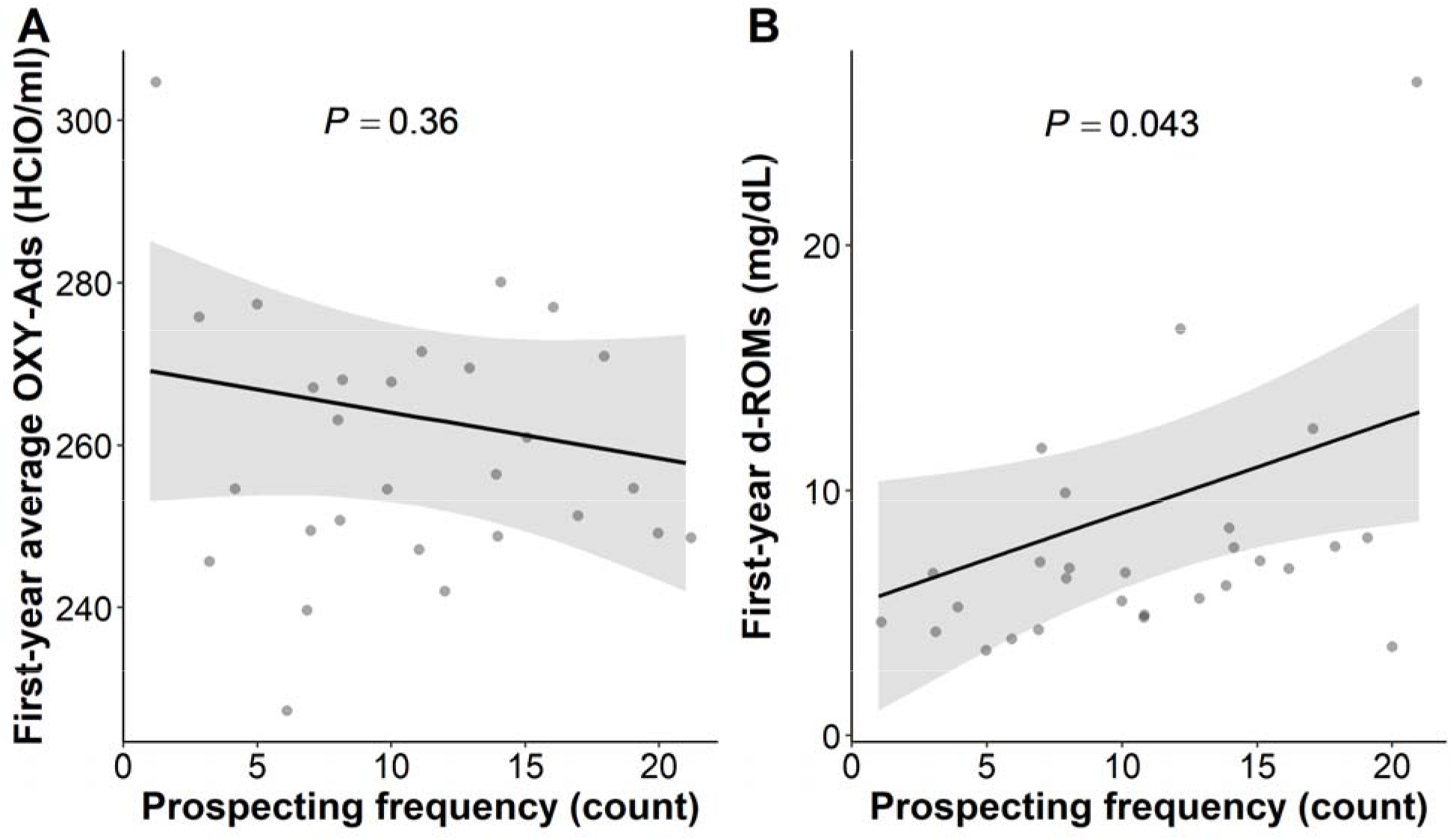
Predicted values (marginal effects) of linear mixed models for specific model terms, (A) first-year average of OXY-Ads and (B) first-year average of d-ROMs, to prospecting frequency (count of observations) for 28 yearling Florida scrub-jays (*N_males_* = 17, *N_females_* = 11). Light gray intervals indicate marginal effects from the models while the dark gray points indicate raw values. Average d-ROMs values are significantly positively correlated with prospecting frequency.

**Table 1.** Linear mixed model results showing relationship between prospecting movement indices and body condition changes throughout the breeding season. Values indicate standardized parameter estimates (β), standard errors (SE), *z*- and *P*-values. Sample sizes for each model is included in parentheses. Bold values indicate significant values with P < 0.05.

| | $\beta$ | SE | $z$ | $P$ |
| --- | --- | --- | --- | --- |
| **OXY-Ads first-year average ($N = 28$)** |  |  |  |  |
| Marginal $R^2 = 0.13$, conditional $R^2 = 0.79$ | | | | |
| **Intercept** | **266.24** | **6.17** | **43.16** | **<0.0001** |
| Prospecting frequency | -3.46 | 3.63 | -0.95 | 0.36 |
| Maximum distance | -7.00 | 5.58 | -1.25 | 0.22 |
| Activity level (sedentary) | -5.28 | 8.03 | -0.66 | 0.52 |
| Sex (male) | -5.55 | 5.02 | -1.11 | 0.30 |
| **OXY-Ads first-year change ($N = 23$)** |  |  |  |  |
| Marginal $R^2 = 0.12$, conditional $R^2 = 0.53$ | | | | |
| **Intercept** | **1.19** | **0.05** | **23.01** | **<0.0001** |
| Prospecting frequency | -0.04 | 0.05 | -0.89 | 0.39 |
| Maximum distance | 0.04 | 0.04 | 1.03 | 0.32 |
| Activity level (sedentary) | 0.02 | 0.08 | 0.19 | 0.85 |
| Sex (male) | -0.03 | 0.06 | -0.61 | 0.55 |
| **d-ROMs first-year average ($N = 28$)** |  |  |  |  |
| Marginal $R^2 = 0.22$, conditional $R^2 = 0.128$ | | | | |
| **Intercept** | **9.24** | **1.73** | **5.34** | **<0.0001** |
| **Prospecting frequency** | **2.30** | **1.07** | **2.15** | **0.04** |
| Maximum distance | -1.08 | 1.24 | -0.87 | 0.41 |
| Activity level (sedentary) | -1.85 | 2.42 | -0.76 | 0.46 |
| Sex (male) | -1.95 | 1.74 | -1.12 | 0.27 |
| **d-ROMs first-year change ($N = 23$)** |  |  |  |  |

| Marginal $R^2 = 0.05$, conditional $R^2 = 0.28$ | | | | |
| --- | --- | --- | --- | --- |
| **Intercept** | **0.94** | **0.29** | **3.21** | **0.005** |
| Prospecting frequency | 0.08 | 0.26 | 0.30 | 0.77 |
| Maximum distance | -0.04 | 0.21 | -0.19 | 0.85 |
| Activity level (sedentary) | -0.23 | 0.46 | -0.50 | 0.62 |
| Sex (male) | 0.19 | 0.32 | 0.58 | 0.57 |
| **Body mass difference during breeding season ($N = 20$)** |  |  |  |  |
| Multiple $R^2 = 0.11$; adjusted $R^2 = -0.11$ | | | | |
| **Intercept** | **0.98** | **0.01** | **106.88** | **<0.001** |
| Prospecting frequency | < -0.001 | 0.01 | -0.07 | 0.95 |
| Maximum distance | -0.01 | 0.01 | -0.73 | 0.47 |
| Activity level (sedentary) | -0.01 | 0.02 | -0.30 | 0.77 |
| Sex (male) | 0.01 | 0.01 | 1.08 | 0.30 |

### Is prospecting condition-dependent?

Early-stage body condition, sex, and age did not influence prospecting effort (*P* > 0.1; Table 2).

**Table 2.** Linear mixed models and generalized linear mixed model results showing relationship between early-stage body condition and subsequent prospecting movement indices for 44 nonbreeding helpers (*N_males_* = 23, *N_females_* = 21) that were observed during aggregation sampling. Values indicate standardized parameter estimates (β), standard errors (SE), *z*- and *P*-values. Bold values indicate significant values with *P* < 0.05, italics indicate P < 0.1.

| | $\beta$ | SE | $z$ | $P$ |
| --- | --- | --- | --- | --- |
| **Prospecting frequency** |  |  |  |  |
| Marginal $R^2 = 0.05$, conditional $R^2 = 0.41$ | | | | |
| **Intercept** | **10.66** | **1.45** | **7.33** | **<0.001** |
| Nestling body condition | 0.11 | 0.87 | 0.13 | 0.90 |
| Juvenile body condition | -0.47 | 0.91 | -0.52 | 0.61 |
| Sex (male) | 1.88 | 1.52 | 1.23 | 0.23 |
| Age (yearling) | -1.54 | 1.62 | -0.95 | 0.35 |
| **Maximum distance** |  |  |  |  |
| Marginal $R^2 = 0.04$, conditional $R^2 = 0.74$ | | | | |
| **Intercept** | **1188.38** | **157.44** | **7.55** | **<0.0001** |
| Nestling body condition | -69.66 | 81.60 | -0.85 | 0.40 |
| Juvenile body condition | -7.74 | 92.57 | -0.08 | 0.93 |
| Sex (male) | -158.90 | 136.64 | -1.16 | 0.26 |
| Age (yearling) | -116.53 | 157.41 | -0.74 | 0.47 |
| **Activity level** |  |  |  |  |
| Marginal $R^2 = 0.19$, conditional $R^2 = 0.33$ | | | | |
| *Intercept* | *-2.05* | *1.02* | *-2.00* | *0.05* |
| Nestling body condition | 0.81 | 0.56 | 1.44 | 0.15 |
| Juvenile body condition | -0.76 | 0.51 | -1.50 | 0.13 |
| Sex (male) | 0.22 | 0.87 | 0.25 | 0.80 |
| Age (yearling) | 0.92 | 0.96 | 0.96 | 0.34 |

### Do early-life morphometrics correspond to oxidative status?

Antioxidant capacity and oxidative damage as yearlings both correlated with nestling body condition and juvenile 7^th^ primary length, but in opposite directions (Figure 5; Table 3). Nestling body condition was negatively correlated with yearling OXY-Ads (β = −10.94, SE = 4.18, *z* = −2.62, *P* = 0.02) but positively correlated with yearling d-ROMs (β = 1.80, SE = 0.32, *z* = 5.59, *P* = 0.007). Juvenile wing length had the opposite effect, with long-winged juveniles having high OXY-Ads (β = 16.96, SE = 4.16, *z* = 4.08, *P* = 0.001) and low d-ROMs (β = −2.40, SE = 0.68, *z* = −3.54, *P* = 0.003). Sex had a significant correlation with yearling OXY-Ads, with females having higher OXY-Ad levels than males (β = −24.03, SE = 8.07, *z* = −2.98, *P* = 0.01). Nestling wing length was correlated with yearling d-ROMs (β = 1.21, SE = 0.48, *z* = 2.53, *P* = 0.049). For yearling OXY-Ads, nestling 7^th^ primary or juvenile body condition had no significant effect (*P* > 0.05). For yearling d-ROMs, juvenile body condition and sex had no significant effect (*P* ≥ 0.1).

**Figure 5.**
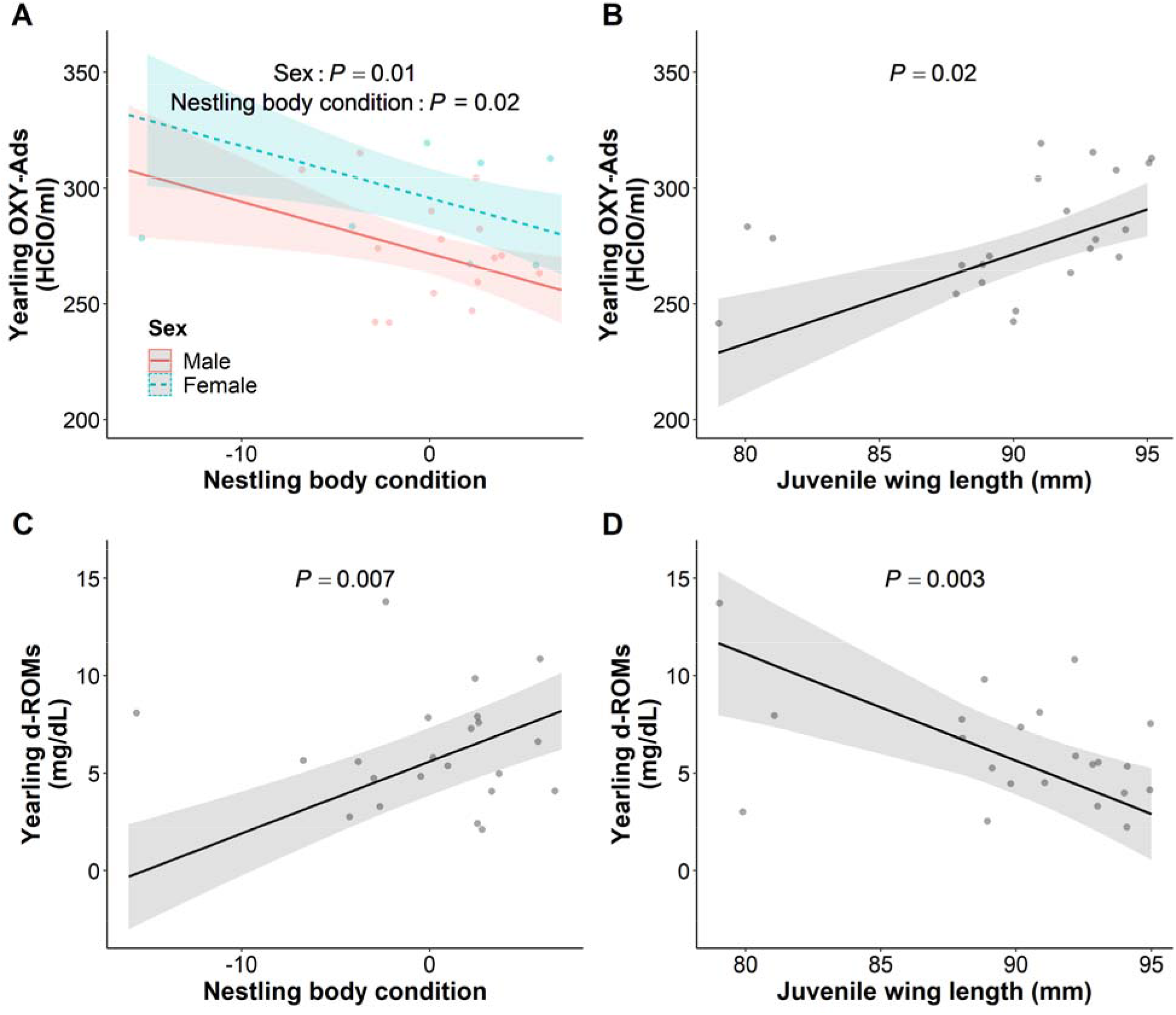
Predicted values (marginal effects) of linear mixed models for specific model terms, (A) yearling OXY-Ads measures and (B) yearling d-ROMs measures for 22 Florida scrub-jays (*N_males_* = 15, *N_females_* = 11). Light gray intervals indicate marginal effects from the models while the dark gray points indicate raw values.

**Table 3.** Linear regression and linear mixed model results showing how early-stage morphometrics and condition affect yearling oxidative status for 22 Florida scrub-jays (*N_males_* = 15, *N_females_* = 11). Values indicate standardized parameter estimates (β), standardized errors (SE), *z*- and *P*-values. Bold values indicate significant values with *P* < 0.05, italics indicate *P* < 0.1.

| | $\beta$ | SE | $z$ | $P$ |
| --- | --- | --- | --- | --- |
| **Yearling OXY-Ads** |  |  |  |  |
| Multiple $R^2 = 0.64$, adjusted $R^2 = 0.53$ | | | | |
| Intercept | **293.80** | **6.44** | **45.63** | **<0.0001** |
| Nestling body condition | **-10.94** | **4.18** | **-2.62** | **0.02** |
| Nestling 7<sup>th</sup> primary | 2.01 | 4.36 | 0.46 | 0.65 |
| *Juvenile body condition* | *6.97* | *3.91* | *1.78* | *0.09* |
| **Juvenile 7<sup>th</sup> primary** | **16.95** | **4.16** | **4.08** | **0.001** |
| Sex (male) | **-24.03** | **8.07** | **-2.98** | **0.01** |
| **Yearling d-ROMs** |  |  |  |  |
| Marginal $R^2 = 0.40$, conditional $R^2 = 0.95$ | | | | |
| Intercept | 5.99 | 0.88 | 6.84 | <0.0001 |
| Nestling body condition | **1.80** | **0.32** | **5.59** | **0.007** |
| **Nestling 7<sup>th</sup> primary** | **1.21** | **0.48** | **2.53** | **0.049** |
| Juvenile body condition | -0.74 | 0.35 | -2.12 | 0.10 |
| **Juvenile 7<sup>th</sup> primary** | **-2.40** | **0.68** | **-3.54** | **0.003** |
| Sex (male) | 0.34 | 0.74 | 0.46 | 0.66 |

## DISCUSSION

While prospecting behaviour precedes and informs natal dispersal, its associated costs such as energetics or increased predation risk have been hypothesised as trade-offs to this information-gathering process (Delgado et al., 2014; Pärt et al., 2011). Our results lend support to this hypothesis, as it appears prospecting movement is associated with certain physiological costs in resident, cooperatively breeding Florida scrub-jays. We found that prospecting frequency correlated with high first-year average oxidative damage. Early-stage body condition did not influence subsequent prospecting when compared across a large set of individuals. However, among our physiologically sampled individuals, early-stage body condition and morphometrics did vary with yearling oxidative status. Nestling body condition was negatively linked to yearling antioxidant capacity while females and long-winged juveniles showed relatively high antioxidant capacity. On the other hand, nestling body condition was positively correlated with yearling oxidative damage, and birds with long wings as nestlings and short wings as juveniles showed high oxidative damage. Overall, our results show that physiological costs to prospecting exist, but individuals do not seem to adjust their prospecting behaviour based on body condition. Rather, the extent of physiological costs seems to depend on early-body condition and morphology, which may explain the observed differences among individuals.

Consistent with our predictions, prospecting frequency was positively correlated with oxidative damage. In homing pigeons, *Columba livia*, those that flew farther distances (200 km) expressed higher oxidative damage and lower antioxidant capacity than those that flew shorter distances (60 km) (Costantini et al., 2008). By comparison, Florida scrub-jays in our population move only relatively short distances during prospecting forays, but those making more frequent movements did show oxidative damage. While we did not detect sex-differences in prospecting movement itself, dispersal is female-biased in Florida scrub-jays with females dispersing earlier and farther than males (Fitzpatrick et al., 1999; Suh et al., 2020). Correspondingly, we observed sex-based differences in oxidative status with females exhibiting higher antioxidant capacity than males as yearlings. Having high antioxidant capacity may be crucial for the dispersing sex, especially as prospecting frequency is linked to oxidative damage. This physiological asymmetry between females and males suggests that prospecting comes at a physiological cost that affects the trade-offs between information acquisition and movement (Delgado et al., 2014; Kokko & Ekman, 2002), shaping individual variation in prospecting effort and subsequent dispersal.

Regarding oxidative status as yearlings, early-life body condition and morphological measures had different effects. Contrary to our prediction, nestling body condition was negatively linked to yearling oxidative status, with nestlings in high body condition exhibiting low antioxidant capacity and high oxidative damage as yearlings. In contrast, juvenile body condition did not affect oxidative status. Given that oxidative measures across life stages are not repeatable within individuals and how morphological development occurs unevenly (Kilgas et al., 2010; Perrig et al., 2014), the ability to combat and deal with oxidative stress seems to also vary depending on developmental stage and condition (Monaghan et al., 2009). For instance, in great tits, *Parus major*, experimentally induced to have hatching asynchrony, late-hatched brood mates exhibited smaller body mass and wing length but not tarsus length or plasma antioxidant potential compared to early-hatched birds (Kilgas et al., 2010). This finding suggests that organisms can balance oxidative damage without compromising development (Hall et al., 2010), or potential outcomes of this trade-off may manifest later in life (Monaghan et al., 2009). Alternatively, parental effects can also influence offspring phenotype. Zebra finches, *Taeniopygia guttata*, that experienced experimentally induced oxidative stress early in life produced fostered offspring with shorter tarsi than those of control fosters that were not manipulated (Romero-Haro & Alonso-Alvarez, 2020). Similarly, our measure of body condition (i.e., residuals from an ordinary least squares linear regression of body mass against tarsus) may have been a product of poor-quality parents producing offspring with short tarsi, in addition to exposing them to elevated oxidative stress from other environmental constraints such as food availability or habitat quality. Indeed, Seychelles warblers from territories with low food availability exhibited higher oxidative damage (van de Crommenacker et al., 2011), suggesting that early-life environmental conditions can affect development mediated by oxidative status. Overall, developmental differences may help explain why individuals vary in subsequent prospecting effort.

In our study, wing length in juveniles was linked to high antioxidant capacity and low oxidative damage as yearlings. However, we cannot yet distinguish cause and effect. On one hand, relatively long flight feathers may simply reflect good oxidative status or associated nutritional status that promoted feather development (Romero et al., 2005). Alternatively, long wings may help juveniles combat oxidative stress as they become mobile and nutritionally independent. For instance, juveniles with longer wings may have better flight capacities, which either prevent depletion of plasma antioxidants (Costantini et al., 2008) or increase access to dietary antioxidants that can mitigate oxidative damage (Catoni et al., 2008; Larcombe et al., 2008). Given the strong correlation between food availability and oxidative damage mentioned above (van de Crommenacker et al., 2011), access to resources enhanced by flight may play a significant role in mediating oxidative status. In addition, evidence shows that birds can mitigate activity-induced oxidative damage by increasing antioxidant-rich food consumption prior to and during migration (Alan & McWilliams, 2013; McWilliams et al., 2021; Skrip & McWilliams, 2016), highlighting the importance of food-related mediation of oxidative stress. We were unable to test this hypothesis in our study.

Curiously, in contrast to juvenile wing length, nestling measures showed opposite relationships with oxidative damage. Again, this effect may have been mediated by oxidative stress affecting development. In red-winged blackbirds, *Agelaius phoeniceus*, individuals provided with dietary antioxidants allocated these extra resources to increasing growth rate, rather than reducing oxidative damage (Hall et al., 2010). While we are unable to determine causal relationships between oxidative status and development, our results highlight the importance of incorporating different measures of body condition to understand individual differences (Labocha & Hayes, 2012; Waye & Mason, 2008).

While oxidative status corresponded to metrics of both movement and morphology, we did not detect body mass changes in prospecting Florida scrub-jays. Previous work on other social vertebrates has yielded examples of body mass loss associated with prospecting efforts (Kingma et al., 2016; Ridley et al., 2008; Young & Monfort, 2009) and even so far as natal dispersal decisions (e.g., Barbraud et al. 2003; Debeffe et al. 2012; but see Loe et al. 2010). We offer several potential explanations for absence of body mass loss in our results. First, contrary to species that effectively float after leaving the natal territory (Maag et al., 2019; Ridley et al., 2008; Young & Monfort, 2009), Florida scrub-jays tend to stay in the vicinity of their natal territory while prospecting and eventually return home (Woolfenden & Fitzpatrick, 1984). This “stay-and-foray” strategy (Reed et al., 1999) may allow individuals to secure and revisit reliable foraging patches (including cached acorns, in the case of Florida scrub-jays) that may be less energetically costly than foraging away from home. Indeed, Florida scrub-jays have year-round caches which may also prevent body mass fluctuations on a daily basis (Lucas, 1994; Pravosudov & Grubb, 1997). It is also possible that constancy of optimal body mass may be critical for flight efficiency in this species (Witter & Cuthill, 1993).

Our results may reflect certain methodological constraints. Measures of oxidative status may be masked by individual regulation of oxidative balance or have long-term consequences that last beyond the scope of this study, such as lifetime reproductive success or survival (Brotons & Broggi, 2003). In Florida scrub-jays, adults manipulated with supplemental corticosterone had reduced chroma in their plumage (Windsor et al., 2019) while juveniles with artificially reduced plumage reflectance lost more dominance interactions (Tringali & Bowman, 2012), indicating complex interactions between physiology, phenotype, and behaviour. Future work involving experimental manipulations and controls on oxidative status may elucidate the relationships between morphological development, body condition, and prospecting efforts, and how these variables ultimately shape lifetime fitness.

Our study is among the first to link multiple repeated measures of body condition with subsequent prospecting effort, demonstrating the complex nature of trade-offs between information-gathering movements and their associated physiological costs. In Florida scrub-jays, prospecting correlated to oxidative damage while early-life conditions and morphology mediated oxidative status later in the lives of pre-breeding individuals. Overall, our results provide insight on the drivers of individual variation in prospecting effort, an understudied life-history behavior having important ramifications to dispersal and ultimately to fitness.

## Notes

### Competing Interest Statement

The authors have declared no competing interest.

