## Supplemental materials for "Physiological costs of prospecting in a resident cooperative breeder, the Florida scrub-jay"

**1. Definition of body condition**

In our study, we used ordinary least squares linear regression of body mass against tarsus as a measure of body condition. However, this estimation of body condition relies on several assumptions that are often not met (Green, 2001). We provide evidence for some of these assumptions: (1) we find that body mass increases linearly with the selected body size indicator (BSI). In our sample, body mass and tarsus length are significantly positively correlated (*P* < 0.005). (2) We also observe that BSI is not correlated with other structural components, namely 7^th^ primary feather length (Table S1).

**Table S1. Correlation matrix between body condition and 7^th^ primary length**

Using Spearman’s correlation coefficient. Top values indicate ρ-values while bottom values indicate *P*-values.

|  | **Nestling body condition** | **Juvenile body condition** | **Nestling 7^th^ primary** | **Juvenile 7^th^ primary** |
| --- | --- | --- | --- | --- |
| **Nestling body condition** |  | 0.19 | 0.29 | 0.26 |
| **Juvenile body condition** | 0.35 |  | 0.35 | 0.08 |
| **Nestling 7^th^ primary** | 0.15 | 0.08 |  | 0.19 |
| **Juvenile 7^th^ primary** | 0.19 | 0.71 | 0.36 |  |

**2. Summary values of prospecting effort and weight change**

**Table S2.** Differences in prospecting effort by sex and age (mean ± SE).

| **Sex** | **Social status** | **N** | **Prospecting frequency** | **Maximum prospecting distance (in m)** |
| --- | --- | --- | --- | --- |
| **Male** | **Yearling** | 13 | 11.08 ± 1.30 | 1115.07 ± 216.17 |
|  | **Older helper** | 19 | 12.84 ± 1.40 | 829.62 ± 127.50 |
| **Female** | **Yearling** | 9 | 9.67 ± 1.83 | 1257.91 ± 276.15 |
|  | **Older helper** | 10 | 11.50 ± 1.94 | 1079.62 ± 208.13 |

**Table S3.** Differences in mean weight loss by sex and age (mean ± SE).

| **Sex** | **Social status** | **N** | **Cumulative weight loss (in g)** |
| --- | --- | --- | --- |
| **Male** | **Yearling** | 13 | -0.39 ± 0.43 |
|  | **Older helper** | 19 | -0.03 ± 0.25 |
| **Female** | **Yearling** | 9 | -1.06 ± 0.65 |
|  | **Older helper** | 10 | 0.83 ± 1.17 |
